# Synergistic Activity of Minocycline and Rifampin in Combination with Antifungal Drugs against *Candida auris*

**DOI:** 10.1101/2021.02.23.432620

**Authors:** Thea Brennan-Krohn, Liam Friar, Sarah Ditelberg, James E. Kirby

## Abstract

*Candida auris* is an emerging multidrug-resistant fungal pathogen that spreads readily in healthcare settings and has caused numerous hospital outbreaks. Very few treatment options exist for *C. auris* infections. We evaluated the activity of all two-drug combinations of three antifungal agents (amphotericin B, caspofungin, and voriconazole) and two antibacterial agents (minocycline and rifampin) against a collection of 10 *C. auris* isolates using an automated, inkjet printer-assisted checkerboard array method. Three antibacterial-antifungal combinations (amphotericin B plus rifampin, amphotericin B plus minocycline, and caspofungin plus minocycline) demonstrated synergistic activity by checkerboard array against ≥90% of strains. The two amphotericin B-containing combinations were also synergistic using the time-kill synergy testing method. Our results suggest that combinations of antifungal and antibacterial agents may provide a promising avenue for treatment of this multidrug-resistant pathogen.

## INTRODUCTION

The pathogenic yeast *Candida auris*, first identified in the external ear canal drainage of a woman in Japan in 2009 (1), was classified by the U.S. Centers for Disease Control and Prevention as one of the most urgent antibiotic resistance threats in 2019 (2). At least 4 distinct clades of the yeast appear to have emerged nearly simultaneously on 3 different continents around the time the first isolate was recognized (3). More recently, invasive *C. auris* infections have been observed as a complication of critical SARS-CoV-2 disease (4–6). *C. auris* is most commonly reported as a cause of bloodstream infections; patients with central venous catheters and recent surgical procedures appear to be at particularly high risk (7). Compared to other *Candida* species, *C. auris* is notable for its propensity to spread rapidly within healthcare settings, for a high rate of mortality in infected patients, and for frequent resistance to multiple antifungal drugs (8, 9).

The number of drugs available to treat even the most susceptible of fungal pathogens is small, with only three classes of systemic antifungal agents in general use: azoles, echinocandins, and the polyene amphotericin B (AMB) (10). Individual patient factors, such as allergies, vulnerabilities to side effects, drug-drug interactions, and the need for penetration into specific tissue sites frequently further constrain the choice of agents that can be safely and effectively used; resistance to any of these agents may reduce practical treatment options to few or none. In this context, the resistance profiles observed among *C. auris* isolates are particularly alarming: nearly all isolates are resistant to fluconazole, and some also demonstrate resistance to echinocandins and AMB (7, 11). Isolates resistant to drugs from all three classes have been reported (12).

Unfortunately, the antifungal drug pipeline is unlikely to yield an abundance of new treatment options in the near future. Antifungal drug development is hampered by intrinsic challenges (fungi, like humans, are eukaryotic organisms, and therefore it is difficult to identify compounds that are active against fungal pathogens but not highly toxic to host cells) and by poor financial incentives for pharmaceutical companies (10). Ibrexafungerp, a first-in-class glucan synthase inhibitor, has recently been shown to have activity against *C. auris* (13) in *in vitro* and animal models (14), but has not yet been approved for clinical use.

One treatment approach that does not rely on the introduction of novel agents is the repurposing of existing drugs in combination. Combination therapy using two or more antifungal drugs is already an established component of therapy for certain fungal infections: AMB is routinely used in combination with 5-fluorocytosine as induction therapy for *Cryptococcus neoformans* meningitis (15), while combinations of antifungal drugs are often employed in an attempt to treat infections caused by highly drug-resistant molds such as *Scedosporium* spp. (16). This strategy, however, still relies on the limited number of currently available antifungal agents.

In 1972, investigators observed *in vitro* synergistic activity in *Saccharomyces cerevisiae* when AMB was combined with rifampin (RIF), an antibacterial RNA synthesis inhibitor (17) or with tetracycline, an antibacterial protein synthesis inhibitor (18); the combination of AMB with RIF was later also noted to be synergistic against several *Candida* species (19). Synergy between AMB and the tetracycline analogue minocycline (MIN) was demonstrated against *Cryptococcus neoformans*, *C. albicans*, and other *Candida* species in 1977 (20). Fluconazole was also subsequently reported to be synergistic with MIN and with another tetracycline analogue, doxycycline, against *C. albicans* (21, 22). To our knowledge, however, these antibacterial-antifungal combinations have not been evaluated in *C. auris*. We used a novel inkjet printer-assisted checkerboard array synergy method as well as time-kill synergy studies to investigate synergistic activity of combinations of MIN, RIF, voriconazole (VRC; an azole antifungal), caspofungin (CAS; an echinocandin), and AMB against a collection of *C. auris* isolates with a range of drug resistance patterns.

## RESULTS

### Single drug MIC testing by broth microdilution (BMD) and digital dispensing method (DDM)

BMD MICs were prepared in triplicate to determine the modal MIC (i.e. the MIC obtained in ≥2 replicates) at 24 hours (all drugs) and 48 hours (VRC and AMB) using standard Clinical and Laboratory Standards Institute (CLSI) methodology (23). If three sequential doubling dilution values were obtained in the three replicates (e.g. 2, 4, and 8 μg/mL), the middle value was considered the modal MIC. Modal MICs ranged from 0.5 to 2 μg/mL for AMB, from ≤0.016 to >8 μg/mL for VRC, and from 0.063-0.5 μg/mL for CAS. VRC MICs were higher at 48 than 24 hours, while AMB MICs were either the same or one doubling dilution higher at 48 hours compared to 24 hours for all strains. Neither RIF nor MIN exhibited inhibitory activity at the concentrations tested (Table 1 and Table S1).

**TABLE 1.** Modal broth microdilution MICs (μg/mL) at 24 and 48 hours

| Strain | AMB | AMB | VRC | VRC | CAS | MIN | RIF |
| --- | --- | --- | --- | --- | --- | --- | --- |
| (AR Bank Number) | 24 hr | 48 hr | 24 hr | 48 hr | 24 hr | 24 hr | 24 hr |
| 0381 | 0.5 | 1 | ≤0.016 | 0.031 | 0.063 | >64 | >128 |
| 0382 | 0.5 | 1 | 0.031 | >8 | 0.125 | >64 | >128 |
| 0383 | 1 | 1 | 1 | 2 | 0.125 | >64 | >128 |
| 0384 | 1 | 1 | 0.5 | N/A | 0.125 | >64 | >128 |
| 0385 | 1 | 1 | 8 | >8 | 0.125 | >64 | >128 |
| 0386 | 1 | 1 | 4 | >8 | 0.125 | >64 | >128 |
| 0387 | 1 | 1 | 0.063 | >8 | 0.125 | >64 | >128 |
| 0388 | 2 | 2 | 1 | 4 | 0.5 | >64 | >128 |
| 0389 | 2 | 2 | 4 | 8 | 0.125 | >64 | >128 |
| 0390 | 1 | 2 | 1 | 2 | 0.125 | >64 | >128 |

AR Bank: CDC & FDA Antibiotic Resistance Isolate Bank; AMB, amphotericin B; CAS, caspofungin; VRC, voriconazole; MIN, minocycline; RIF, rifampin. N/A: Modal MIC could not be determined as MICs were 0.125, 2, and 8 µg/mL

For each condition (i.e., each drug tested against each strain at 24 or 48 hours), five DDM MIC values were also obtained: a dedicated MIC test result and four results from the single-drug titrations of each synergy grid in the checkerboard array testing described below (Table S1). When on-scale DDM MIC results were compared to on-scale modal BMD MICs, 94.7% were within ±1 doubling dilution and 99.5% were within ±2 doubling dilutions of the modal MIC (Table 2). In antifungal susceptibility testing, essential agreement between a new method and the reference method is generally defined as an MIC value falling within ±2 doubling dilutions of the reference result (24, 25). Concordance with the modal BMD MIC was highest for AMB (100% of on-scale results within ±1 doubling dilution) and lowest for CAS (81.6% and 98.0% of on-scale results within ±1 and ±2 doubling dilutions, respectively). Therefore, our data indicate that the DDM automation method provides accurate and robust antifungal testing results.

**TABLE 2.** Essential agreement between on-scale DDM MIC results and modal BMD MIC

| Drug and time | DDM MIC results within $\pm 1$ two-fold | DDM MIC results within $\pm 2$ two-fold |
| --- | --- | --- |
|  | dilution of modal MIC (*n*, %) | dilutions of modal MIC (*n*, %) |
| AMB, 24 hr | 50/50 (100.0) | 50/50 (100.0) |
| AMB, 48 hr | 49/49 (100.0) | 49/49 (100.0) |
| CAS, 24 hr | 40/49 (81.6) | 48/49 (98.0) |
| VRC, 24 hr | 35/37 (94.6) | 37/37 (100.0) |
| VRC, 48 hr | 21/21 (100.0) | 21/21 (100.0) |
| MIN, 24 hr | N/A | N/A |
| RIF, 24 hr | N/A | N/A |
| Total | 195/206 (94.7) | 205/206 (99.5) |

AMB, amphotericin B; CAS, caspofungin; VRC, voriconazole; MIN, minocycline; RIF, rifampin.
MIC results were not used when >1 skipped well occurred (*n* = 6), when modal BMD MIC or DDM result was off-scale (*n* = 33), or when there was no modal BMD MIC (*n* = 5).
N/A: Essential agreement could not be calculated due to off-scale high modal BMD MICs; all DDM results were also off-scale high for these drugs.

### Checkerboard array synergy testing using DDM

Using the checkerboard array assay, we found that MIN was synergistic against all 10 strains when combined with either AMB or CAS, and RIF was synergistic with AMB against 9 strains. In synergistic combinations, the concentration of MIN at the FIC_I-MIN_ ranged from 4-16 μg/mL and was ≤8 μg/mL in 12 of 22 cases, while the concentration of RIF was 8 μg/mL in 5 cases and 16 μg/mL in 4 cases. All other combinations were synergistic against ≤2 strains. Three of the 5 combinations that were not synergistic against any strains were antagonistic against 2 strains. (Table 3 and Table S2).

**TABLE 3.**
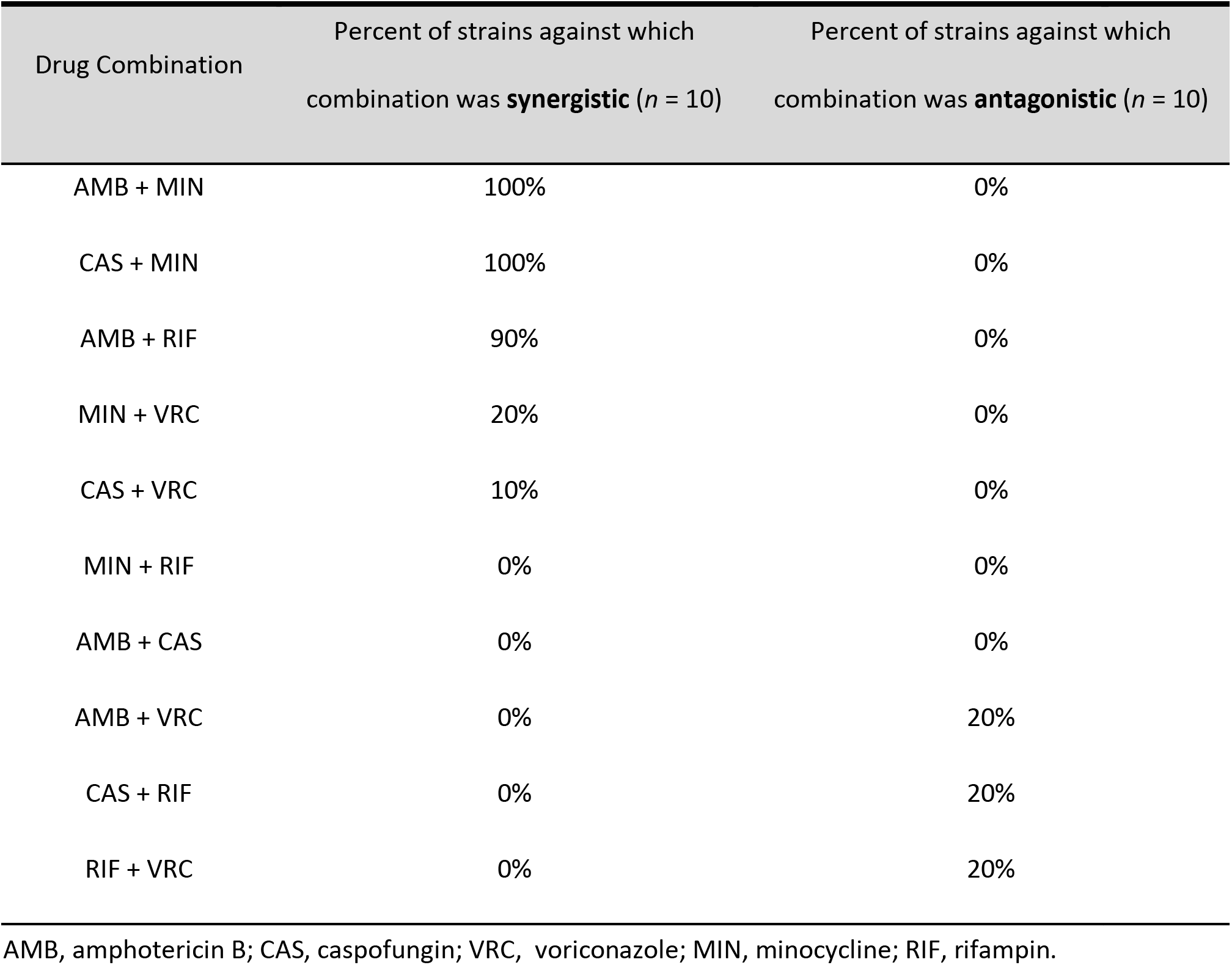
Checkerboard array synergy results

### Time-kill synergy testing

The three combinations that demonstrated synergy against ≥9 strains by the checkerboard array assay (MIN plus AMB, MIN plus CAS, and RIF plus AMB) were tested using the time-kill method against a subset of 5 of the *C. auris* strains, which were chosen to represent a range of susceptibility profiles. (Table 4, Figure 1). Each drug pair was tested, at two combinations of concentrations of individual drugs, against each of the 5 isolates, and was considered synergistic and/or fungicidal if at least one of these combinations met criteria for synergistic or fungicidal activity, respectively. At 24 hours, AMB plus RIF and AMB plus MIN were synergistic against all 5 strains, while at 48 hours, AMB plus RIF was synergistic against 4 strains and AMB plus MIN against 3 strains. In combination with RIF, AMB showed fungicidal activity against 2 strains at 24 and 48 hours, and against an additional strain at 24 hours, at concentrations at which it was not fungicidal alone. In combination with MIN, AMB showed fungicidal activity against 2 strains at 24 and 48 hours at concentrations at which it was not fungicidal alone. The combination of CAS plus MIN was not synergistic or fungicidal against any strains. The synergy killing curves for this combination were notable in that CAS exhibited an inhibitory or near-inhibitory effect even at concentrations at or below the BMD MIC, yet its effect was unchanged by the addition of MIN. To further investigate this finding, single-drug killing curve studies were performed with CAS over a range of concentrations; these showed minimal effect of drug concentration on inhibition or killing, in accordance with previously reported observations (26) (Figure 2).

**TABLE 4.** Time-kill results

| Strain<br>(AR Bank<br>Number) | Drug concentrations<br>($\mu\text{g/mL}$) | Concentration difference of cells treated<br>with combination vs most active single agent<br>($\log_{10}$ CFU/mL) | | Concentration difference at end vs start of<br>cells treated with combination<br>($\log_{10}$ CFU/mL) | |
| --- | --- | --- | --- | --- | --- |
|  |  | Shaded cells indicate *synergy* |  | Shaded cells indicate *fungicidal activity* |  |
|  |  | 24 hours | 48 hours | 24 hours | 48 hours |
| 0381 | AMB 0.125 + RIF 4 | -4.14 | -2.17 | -2.94 | -0.61 |
|  | AMB 0.25 + RIF 4 | -2.03 | -4.30 | -3.12 | -3.12 |
| 0382 | AMB 0.25 + RIF 4 | -2.37 | -4.78 | -3.21 | -3.21 |
|  | AMB 0.5 + RIF 4 | 0.00 | -2.85 | -2.70 | -2.70 |
| 0383 | AMB 0.125 + RIF 8 | -1.80 | -0.50 | 0.44 | 1.74 |
|  | AMB 0.25 + RIF 8 | -3.98 | -3.83 | -2.39 | -1.99 |
| 0388 | AMB 0.25 + RIF 16 | -0.40 | 0.07 | 1.57 | 1.91 |
|  | AMB 0.5 + RIF 16 | -4.82 | -0.82 | -2.78 | 1.05 |
| 0389 | AMB 0.5 + RIF 16 | -3.04 | -0.80 | -1.38 | 1.00 |
|  | AMB 1 + RIF 16 | -2.43 | -1.51 | -3.52 | -0.30 |
| 0381 | AMB 0.125 + MIN 8 | -2.30 | -0.60 | -1.15 | 0.95 |
|  | AMB 0.25 + MIN 8 | -2.08 | -4.78 | -3.19 | -3.19 |
| 0382 | AMB 0.125 + MIN 8 | -2.34 | -0.75 | -0.80 | 1.41 |
|  | AMB 0.25 + MIN 8 | -1.30 | -2.76 | -1.64 | -1.16 |
| 0383 | AMB 0.125 + MIN 8 | -2.02 | -0.01 | 0.13 | 2.02 |
|  | AMB 0.25 + MIN 8 | -3.12 | -2.34 | -1.30 | -0.60 |
| 0388 | AMB 0.5 + MIN 8 | -2.37 | -0.06 | -0.60 | 1.74 |
|  | AMB 1 + MIN 8 | 0.62 | -1.30 | -0.64 | -0.43 |
| 0389 | AMB 0.5 + MIN 16 | -0.53 | 0.11 | -0.96 | 1.77 |
|  | AMB 1 + MIN 16 | -3.48 | -4.99 | -3.30 | -3.30 |
| 0381 | CAS 0.031 + MIN 8 | -0.34 | -0.34 | -0.02 | -0.18 |
|  | CAS 0.031 + MIN 16 | -0.46 | 0.00 | -2.46 | -2.00 |
| 0382 | CAS 0.125 + MIN 8 | 1.07 | 0.84 | -0.27 | -0.36 |
|  | CAS 0.125 + MIN 16 | 1.02 | 0.96 | -1.24 | -1.78 |
| 0383 | CAS 0.063 + MIN 4 | 0.65 | 1.26 | 0.20 | 0.34 |
|  | CAS 0.063 + MIN 8 | 0.63 | 1.00 | -1.81 | -1.57 |
| 0388 | CAS 0.125 + MIN 8 | 0.35 | 0.85 | -0.18 | 0.44 |
|  | CAS 0.125 + MIN 16 | 0.27 | 0.39 | -1.66 | -1.55 |
| 0389 | CAS 0.25 + MIN 1 | 0.77 | 0.65 | 0.37 | 0.20 |
|  | CAS 0.25 + MIN 2 | 0.38 | 0.78 | -1.72 | -1.46 |

AR Bank: CDC & FDA Antibiotic Resistance Isolate Bank; AMB, amphotericin B; CAS, caspofungin; MIN, minocycline; RIF, rifampin.

**FIG 1.**
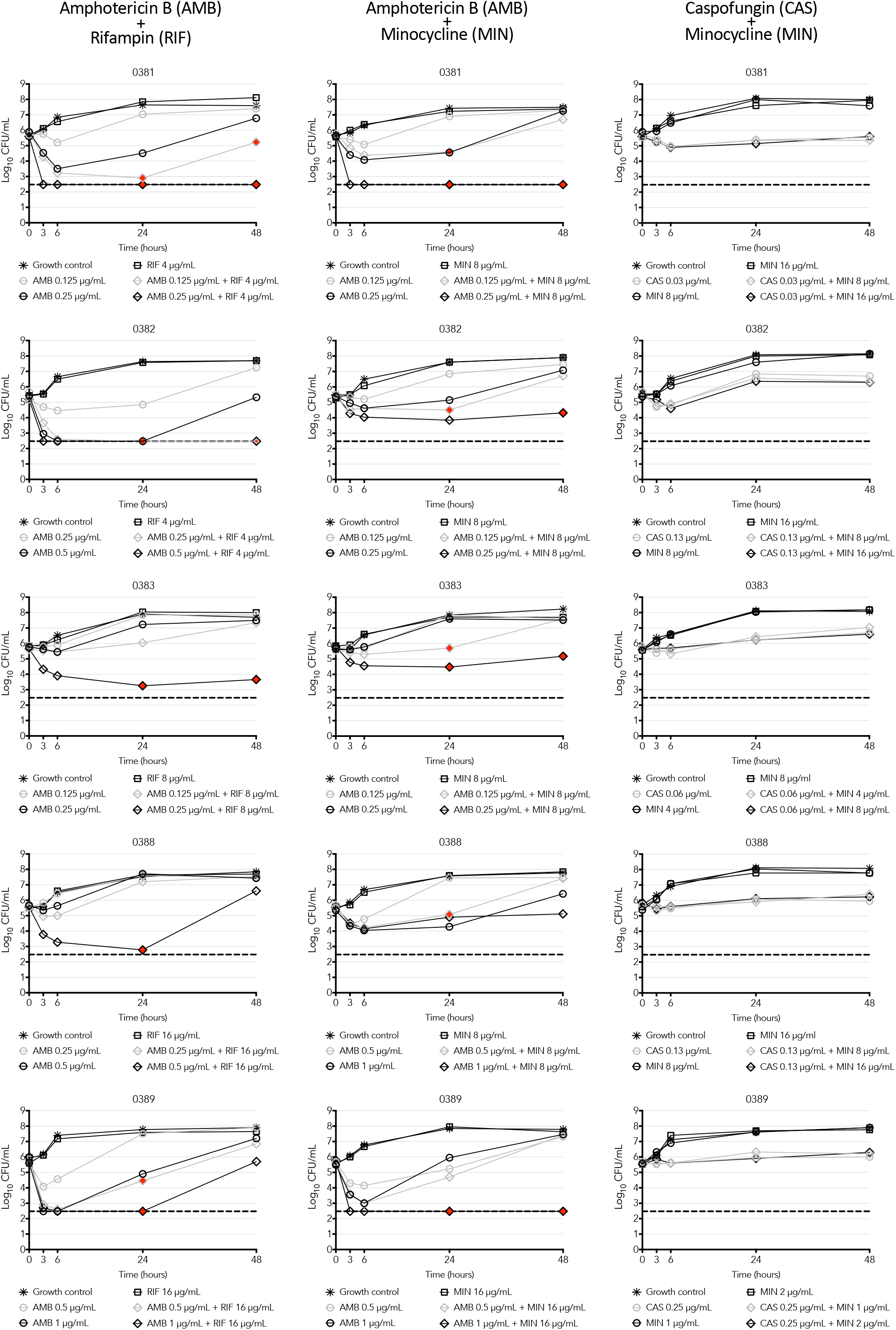
Time-kill synergy graphs. Strain numbers refer to CDC & FDA Antibiotic Resistance Isolate Bank designations. Dashed line indicates assay lower limit of detection. Filled (red) symbols indicate synergistic concentration combinations.

**FIG 2.**
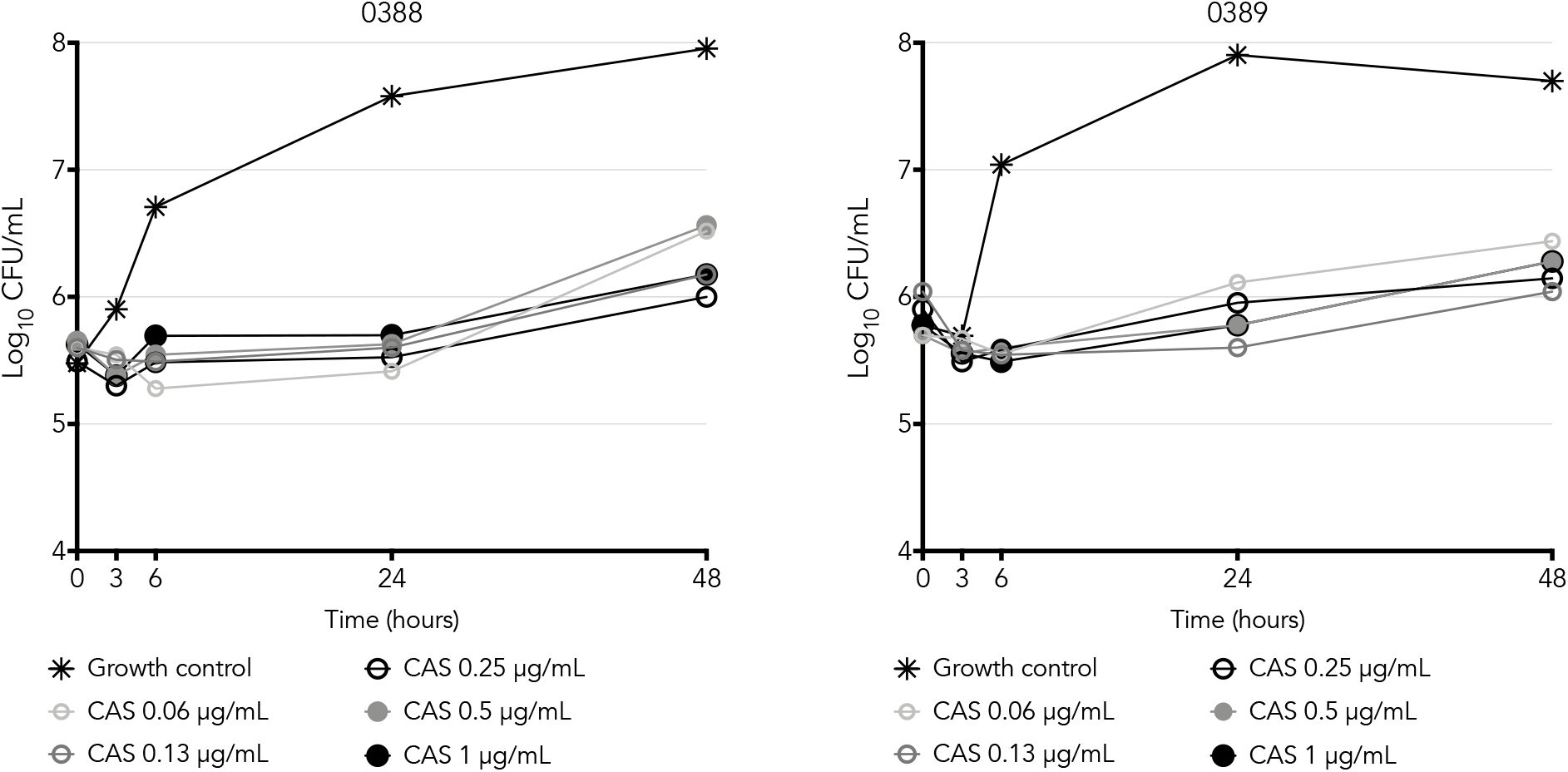
Caspofungin killing curves. Strain numbers refer to CDC & FDA Antibiotic Resistance Isolate Bank designations. The broth microdilution MIC of caspofungin is 0.5 μg/mL for strain 0388 and 0.13 μg/mL for strain 0389.

## DISCUSSION

We identified three combinations of antibacterial and antifungal drugs (AMB plus RIF, AMB plus MIN, and CAS plus MIN) that demonstrated synergistic activity by checkerboard array against ≥90% of *C. auris* strains evaluated; the two AMB-containing combinations were also synergistic by time-kill synergy testing at one or more concentration combinations against the strains evaluated. We thereby demonstrate synergistic activity between antibacterial and antifungal agents against the emerging and highly multidrug-resistant pathogen *C. auris*.

We hypothesize that the mechanism of synergy for these combinations involves impairment in cell wall or membrane integrity by the antifungal drug, permitting entry of an antibacterial agent that would otherwise be unable to access the intracellular compartment of a fungal cell. Echinocandins such as CAS act by inhibiting β-(1,3)-glucan synthase, thus impeding cell wall synthesis and impairing cell wall integrity, resulting in increased vulnerability of the cell to osmotic pressure (27). AMB has traditionally been understood to act by forming ion channels in the lipid bilayer of the fungal cell membrane, thus permeabilizing the cell and ultimately causing cell death (28, 29). However, recent work suggests that large aggregates of AMB, which assemble outside the membrane and act as “sterol sponges” that kill cells by extracting ergosterol from the lipid bilayer, may play a more important role in cell death than do the ion channels (30). It is conceivable that at the subinhibitory concentrations at which it demonstrates synergy with MIN and RIF, AMB could be exerting ion channel permeabilizing activity without aggregate-based cytotoxicity, allowing entry of the antibacterial drugs through the channels.

Our hypothesis is supported by prior observations that tetracycline inhibits protein synthesis in isolated yeast ribosomes (31) and that RIF appears to inhibit RNA polymerase in yeast (17). These drugs, therefore, may have targets in yeast cells analogous to those in bacteria but accessible in yeast only in the setting of disruption of cell membrane or cell wall integrity. A similar phenomenon is well established in bacteria, whereby drugs that are unable to bypass the defenses of the Gram-negative outer membrane under normal circumstances demonstrate activity in the presence of low levels of membrane-permeabilizing agents such as polymyxins (32, 33).

We used two separate synergy testing methods in an effort to increase the robustness of our results, and found that while two combinations demonstrated consistent synergy using both methods, a third was synergistic only by the checkerboard array method and not the time-kill method. It is possible that this finding reflects a limitation of the synergistic activity of this combination (CAS plus MIN). Alternatively, the finding may reflect the limitations of the time-kill testing method for evaluation of CAS activity, as our single-drug CAS killing curves did not demonstrate concentration-dependent inhibitory or fungicidal effects, a finding that has been previously observed with this drug in *C. auris* (26).

In this study we also demonstrated the utility of an automated inkjet printer-assisted digital dispensing method for MIC and checkerboard array synergy testing in yeast. Manual synergy testing is an error-prone and time-consuming process, and automation allows for significantly higher throughput of the technique, thereby facilitating more rapid investigation of novel combinations. This use of the DDM for MIC and synergy testing of bacteria, first described in our laboratory (34, 35), has been adopted by the United States Centers for Disease Control and Prevention to test drug combinations against multidrug-resistant bacterial pathogens (36) and has the potential for similar use for fungal pathogens such as *C. auris*.

*In vitro* synergy testing has certain intrinsic limitations and is not always a direct indicator of *in vivo* efficacy (37). However, identification of combinations with *in vitro* activity provides preliminary data to suggest regimens that may ultimately prove to be of therapeutic benefit. Given the paucity of new antifungal drugs in the development pipeline, regimens that involve readily available drugs for which extensive pharmacokinetic and safety data already exist offer the potential for expedited clinical evaluation and implementation. The concentrations of antifungal drugs active in the combinations identified were very low and easily clinically achievable. Although no interpretive criteria (i.e. susceptibility breakpoints) exist for MIN or RIF for yeast, the concentration of MIN in synergistic combinations was ≤8 μg/mL in more than half the instances of synergy we identified; such concentrations would be considered susceptible (4 μg/mL) or intermediate (8 μg/mL) for Gram-positive bacterial pathogens such as *Staphylococcus aureus* and *Enterococcus* spp. by CLSI (38), suggesting plausible clinical applicability. In topical or local applications (e.g. ophthalmic drops or catheter coating), antibiotics can often be used at concentrations greater than can be safely achieved systemically (39); the combinations we identified could thus also have potential use in these scenarios.

The need for new therapeutic options for *C. auris* has been underscored since the advent of the COVID-19 pandemic, with recent reports from India (4), Colombia (5), and the United States (6) describing *C. auris* infection as a complication of SARS-CoV-2-related critical illness. In addition to possible direct applicability, if the combinations we evaluated act, as we predict, by allowing access of MIN and RIF to intracellular targets in yeast, this information may guide future antifungal drug development approaches. Evaluation of combinations in animal models and ultimately in clinical trials will be critical future steps in establishing clinical activity. In the absence of a predictable timeline for introduction of novel antifungal agents, repurposing existing drugs may be our best hope in identifying new treatment approaches for patients with infections caused by *C. auris* and other emerging multidrug-resistant fungal pathogens.

## MATERIALS AND METHODS

### Fungal isolates

Ten *C. auris* isolates were obtained from the CDC & FDA Antibiotic Resistance (AR) Isolate Bank (Atlanta, GA). *Candida parapsilosis* ATCC 22019, *Candida krusei* ATCC 6258, *Escherichia coli* ATCC 25922, and *Staphylococcus aureus* ATCC 29213 were obtained from the American Type Culture Collection (Manassas, VA). All strains were colony purified, minimally passaged, and stored at −80°C in tryptic soy broth (BD Diagnostics, Franklin Lakes, NJ) with 50% glycerol (Sigma-Aldrich, St. Louis, MO).

### Antimicrobial agents

Voriconazole (VRC) was obtained from Acros Organics (Pittsburgh, PA). Caspofungin (CAS) was obtained from Carbosynth (Oakbrook Terrace, IL). Amphotericin B (AMB) was obtained from Sigma-Aldrich (St. Louis, MO). Minocycline (MIN) was obtained from Chem Impex International (Wood Dale, IL). Rifampin (RIF) was obtained from Fisher Scientific (Waltham, MA). Antimicrobial stocks were prepared in DMSO (Sigma-Aldrich), with the exception of minocycline stock used for time-kill experiments, which was prepared in water. All antimicrobials were quality control (QC) tested with *C. parapsilosis* ATCC 22019 and *C. krusei* ATCC 6258 (VRC, CAS, and AMB) or with *E. coli* ATCC 25922 and *S. aureus* ATCC 29213 (MIN and RIF) and were used only if they produced an MIC result in the QC range accepted by CLSI (38, 40). After passing QC, stocks were aliquoted and stored at −80°C (antifungal drugs) or −20°C (antibacterial drugs) until use. Aliquots were discarded after a single use.

### Antimicrobial susceptibility testing

Manual broth microdilution (BMD) testing of *C. auris* isolates was performed in triplicate for each drug according to CLSI guidelines (23). Strains were isolation streaked on Sabouraud dextrose agar plates (Thermo Scientific, Waltham, MA) and incubated for 24 hours at 35°C in ambient air. BMD plates were made by preparing serial 2-fold dilutions of antimicrobial agents at twice the desired final concentration in 100 μL RPMI 1640 media with L-glutamine (Cytiva, Marlborough, MA) prepared with MOPS buffer (Fisher Scientific, Waltham, MA) in clear, round-bottom, untreated 96-well plates (Evergreen Scientific, Los Angeles, CA, USA). Fungal inocula were prepared by suspending colonies from the overnight plates in sterile 0.9% sodium chloride and adjusting to a 0.5 McFarland standard, diluting this suspension 1:1000 in RPMI, then adding 100 μL of the diluted suspension to each well for a final volume of 200 μL and cell density of 0.5-2.5 ×10^3^ CFU/mL. Negative (sterility) control and growth control wells were included in each row. Plates were then incubated at 35°C in ambient air. At 24 hours (all drugs) and 48 hours (VRC and AMB), plates were removed from the incubator and vortexed on a plate shaker for 4 minutes, after which OD_600_ readings were taken with a Tecan Infinite M1000 Pro microplate reader (Tecan, Morrisville, NC) to quantify growth. OD_600_ readings were normalized by subtracting the average reading of the negative control wells from the same plate, which contained media without yeast, then the percent inhibition for each well was calculated relative to the average of the positive growth control wells of the same isolate from the same plate. For CAS and VRC, the lowest concentration of drug that reduced growth by at least 50% was considered the MIC; for AMB the lowest concentration of drug that reduced growth by at least 90% was considered the MIC (23). If a skipped well occurred, MIC testing was repeated.

MIC testing was also performed using an automated inkjet printer-assisted digital dispensing method (DDM) adapted from a method developed in our laboratory for MIC testing of bacteria (34). Initial 0.5 McFarland yeast suspensions were prepared in RPMI as described above and then diluted 1:2000 in RPMI. The same final volume and cell density in each well was achieved by adding 200 μL of this diluted suspension to each well in a 96-well plate. Antimicrobial drugs were then dispensed by the HP D300 digital dispenser instrument (HP, Inc., Palo Alto, CA) into the yeast suspension in the wells. Incubation and growth interpretation were carried out as in the BMD method described above.

### Checkerboard array synergy testing

The DDM method described above was used to prepare checkerboard arrays in which two drugs were each dispensed in 7-9 two-fold dilutions. Each combination was tested against every *C. auris* strain, with growth determinations made after 24 hours of incubation. Growth inhibition was determined as described above. For combinations in which both drugs use 90% inhibition for MIC determination (AMB plus MIN, AMB plus RIF, and MIN plus RIF), combination wells were considered inhibitory when growth was inhibited by 90% relative to growth control wells; for all other combinations, combination wells were considered inhibitory when growth was inhibited by at least 50% relative to growth control wells. For each combination well in which growth was inhibited, the fractional inhibitory concentration (FIC) for each drug was calculated by dividing the concentration of the drug in that well by the MIC of that drug alone. The FIC index (FIC_I_) for the well was then calculated by summing the FICs of the two drugs in the well; in cases where the MIC of a drug was off-scale, the highest concentration tested was assigned an FIC of 0.5 to permit calculation of the FIC_I_. A combination was considered synergistic against an isolate if it had a minimum FIC_I_ (FIC_I-MIN_) of ≤0.5, antagonistic if it had an FIC_I-MIN_ of >4.0, and indifferent if it had an intermediate FIC_I-MIN_ value (41).

### Time-kill synergy testing

Antimicrobial stocks were diluted in 9.5 mL of RPMI 1640 in 25-by 150-mm glass round-bottom tubes to the appropriate starting concentrations, which were selected based on checkerboard array synergy results. For the AMB-containing combinations, two different AMB concentrations (chosen based on checkerboard array results) were tested in combination with a fixed concentration of MIN or RIF. For the combination of CAS plus MIN, two concentrations of MIN were tested with a fixed concentration of CAS because of the observation that the effect of CAS on yeast growth was minimally affected by CAS concentration (see results and discussion). Negative (sterility) control and positive growth control tubes containing no antimicrobials were also prepared. A 1.0 McFarland suspension of yeast cells from an overnight plate was prepared in 0.9% sodium chloride and 0.5 mL of this suspension was added to each tube for a final starting concentration of 1-5×10^5^ CFU/mL. Cultures were incubated with shaking in ambient air at 35°C for 48 hours. At 0, 3, 6, 24, and 48 hours, aliquots were removed from the culture tube and a 10-fold dilution series was prepared in 0.9% sodium chloride. A 10 μL drop from each dilution was plated onto Sabouraud dextrose agar and incubated overnight. The colonies within each drop were then counted; drops containing 3 to 40 colonies were considered usable and cell density was calculated from these. If more than one dilution for a given sample was usable, the cell densities of the two drops were averaged. If no drops were usable, the densities for consecutive drops above and below the usable range were averaged. The lower limit of detection with this method is 300 CFU/mL. A combination was considered synergistic if it resulted in a ≥2 log_10_ reduction in CFU/mL compared to the most active agent alone and fungicidal if it resulted in a ≥3 log_10_ CFU/mL reduction compared to starting inoculum. Synergy and fungicidal activity were evaluated at 24 and 48 hours.

### Data analysis

Data output from plate readings was visualized using Microsoft Excel (Microsoft Corporation, Redmond, WA). A custom Python script was used to normalize MIC and synergy results and to calculate and visualize growth inhibition.

## ACKNOWLEDGEMENTS

T.B.-K. was supported by a Eunice Kennedy Shriver National Institute of Child Health and Human Development pediatric infectious diseases research training grant (T32HD055148), a National Institute of Allergy and Infectious Diseases (NIAID) training grant (T32AI007061), a Boston Children’s Hospital Office of Faculty Development Faculty Career Development fellowship, an Academy of Clinical Laboratory Physicians and Scientists (ACLPS) Paul E. Strandjord Young Investigator Grant, and a NIAID career development award (1K08AI132716). The HP D300 digital dispenser and TECAN M1000 used in experiments were provided by TECAN (Morrisville, NC). TECAN had no role in study design, data collection/interpretation, manuscript preparation, or decision to publish.

